# Impact of acute chlorpyrifos toxicity on the histology and biochemistry of the stomach, intestine, and muscle of the *Anabas testudineus*

**DOI:** 10.1101/2023.10.29.564571

**Authors:** Bindu Vijayakumari Sudhakaran, S Sreeja

**Author notes:** Email id (Bindu Vijayakumari Sudhakaran).

## Abstract

Chlorpyrifos is classified as an organophosphorus pesticide that exhibits properties of being a broad-spectrum insecticide, acaricide, and nematicide. It is characterized by its ability to degrade quickly and its non-persistent nature. The current investigation involved the determination of the LC_50_ value of chlorpyrifos on *A. testudineus*. The computed LC_50_ values were found to be 11.59, 8.82, 6.30, and 3.82 ppm for exposure durations of 24, 48, 72, and 96 hours, respectively. The levels of protein and glycogen in the stomach, gut, and muscle of A. testudineus, which were subjected to a concentration of 3.82 ppm (LC_50_ 96 hours) chlorpyrifos for a duration of 96 hours, exhibited a notable reduction when compared to the control group. The stomach sections of A. testudineus, when subjected to a concentration of 3.82 ppm chlorpyrifos for a duration of 96 hours, exhibited degeneration of the mucosa layer and necrosis of the mucosa epithelium. The histopathological examination of the intestine revealed several notable findings, including the presence of a disintegrating serosa, a significantly enlarged muscularis layer, and observable damage to both the circular muscle layer and submucosa. Additionally, evidence of necrosis was observed within the mucosa layer. The muscle tissue exhibited degeneration inside the muscle bundles, characterized by the presence of discrete regions of necrosis and the development of vacuoles within the muscle bundles. The results of this study demonstrate that chlorpyrifos exhibits a significant level of toxicity towards A. testudineus, and it is important to note that the absence of mortality does not guarantee the maintenance of physiological well-being in this particular fish species.

## I. Introduction

The utilization and advancement of pesticides to manage a diverse range of herbaceous and insectivorous pests, which can negatively impact the production of food in terms of both quality and quantity, have played a pivotal role in the Green Revolution. This transformative movement, which has reshaped society since the 1950s, aligns with the era commonly referred to as the “chemical age”. In specific instances, the advantages of chemistry can also manifest as disadvantages, since they contribute to the decline of biodiversity and pose a threat to the viability of significant ecosystems by disrupting predator-prey dynamics. Agriculture is widely acknowledged as a crucial sector in India, serving as the foundation of the country’s economy. However, it is worth noting that a significant portion, approximately 70%, of the insecticides and herbicides employed in agricultural practices are synthetic chemicals. These chemicals have been found to have detrimental effects on the health of farmers, primarily through inhalation or dermal penetration during the spraying process. Such adverse effects may manifest through skin lesions or wounds. These pollutants are introduced into the aquatic ecosystem through the process of leaching into groundwater tables and surrounding water bodies. Through the substitution of the enduring and challenging organochlorine compounds Organophosphates (OPs) have emerged as the predominant class of insecticides on a global scale. Assessing the toxicity of these pesticides on the ecosystem presents challenges due to their limited solubility, quick breakdown, and brief stability in the water column. Chlorpyrifos is classified as an organophosphorus pesticide, possessing properties as a broad-spectrum insecticide, acaricide, and nematicide. It is chemically identified as O,O-diethyl O-(3,5,6-trichloro-2-pyridyl) phosphorothioate. The substance in question is a solid with a color ranging from amber to white, possessing a crystalline structure (Mackay et al., 1999). It finds extensive application in India for many objectives, including indoor pest control, protection of various food crops, management of structural pests, and maintenance of grass and decorative plants (NPIC, 2017). The nerve tissue is the specific target organ of pests for Chlorpyrifos. Similar to other organophosphorus pesticides, Chlorpyrifos acts by inhibiting the enzyme acetylcholinesterase, leading to its conversion into choline and acetate (Eaton et al., 2008).

Fish are well recognized as effective bioindicators of the aquatic environment due to their ability to offer significant insights into the overall health and ecological stability of the aquatic ecosystem (Bartoskova et al., 2013; Faggio et al., 2014a, b; Gobi et al., 2018).Various fish species have been found to experience hepatic dysfunction, neurobehavioral and neurochemical alterations, genotoxicity, and lesions in their gills, eyes, and brain with exposure to chlorpyrifos, as described by Cengiz and Unlu (2006), Deb and Das (2013), and Zahran et al. (2018).Chemical formulations employed in agricultural practices to manage pests and improve crop yields have the potential to infiltrate various ecological compartments, including rivers, lakes, and ponds, via leaching, agricultural runoff, and wind dispersion. Consequently, these chemicals can exert adverse effects on numerous organisms and ecosystems that extend beyond their intended target (Bhatnagar and Bana, 1992). Ensuring the well-being of fish is of paramount significance due to the substantial contribution of fish sources to the global food supply (Tripathi and Harsh 2002).The species Anabas testudineus, commonly known as the climbing perch, is a freshwater fish that is consumed as food in the state of Kerala, India. The objective of this research was to assess the potential toxicity of chlorpyrifos, an insecticide, on this particular species of hardy freshwater fish.

## II. MATERIALS AND METHODS

Quantification of the potential exposure of pesticides, in the form of concentrations in the exposure media that may contact the human body is needed for the direct exposure estimation of pesticides and quantification, follows from exposure data. This modeling involves quantification of the contact with the exposure media containing the substance by defining exposure routes and exposure patterns, including contact durations, contact frequencies and site of contact. In the present study, hardy edible freshwater fish *Anabas testudineus* was selected as the test organism. Fishes contribute the main source of protein in our diet and also the health of the fish reflects the health status of the aquatic environment.

### Test organism-Anabas testudineus

An organism selected for a toxicity test is mainly based on certain criteria like its position within the food chain, suitability for laboratory studies, ecological status, genetic stability, uniform populations and adequate background data on the organism. For the present study freshwater teleost *Anabas testudineus* was selected due to its wide availability and suitability as a model for toxicity testing and also due to sustainability in laboratory conditions. With the changing environment,*Anabas testudineus* shows a well-adaptive nature.

### Collection and handling of the test animal

Uniform sized air-breathing fish, Anabas testudineus (B1och), was procured from a nearby stocking pond. The test organisms were transported in big circular plastic containers in order to mitigate the risk of overcrowding and self-inflicted harm caused by collisions with the container walls. During transportation to the laboratory, measures were taken to ensure that the test organisms were not subjected to overcrowding, bruising, or stress, which consequently reduced their vulnerability to disease. In order to prevent cutaneous infection, the subjects were subjected to a five-minute treatment with a 0.1% solution of potassium permanganate, resulting in disinfection.

The fish underwent a 15-day acclimation period, during which only individuals displaying signs of activity and good health were chosen for further acclimation. During the acclimatization period, the subjects were provided with laboratory-prepared food consisting of rice bran, tapioca, fish meal, ground nut, oil cake, and a sufficient quantity of vitamins. The feeding occurred on a daily basis, with 2-3 meals provided each day. In all toxicity trials, animals that were healthy and exhibited normal activity levels, and were approximately of equal size, were randomly chosen from the holding tanks and employed. studies were conducted using fish specimens measuring 14-15 cm in total length and weighing 45-50g. Prior to the studies, the fish were subjected to a 24-hour period of starvation, during which their gastrointestinal tracts were completely emptied of all ingested food and waste materials.

### Parameters of Water Quality

The physio-chemical properties of water were measured, including temperature (27.15 ± 2.51°C), pH (7.16 ± 0.05), dissolved oxygen (7.49 ± 0.31 mg/L), and free carbon dioxide (6.51 ± 0.9 mg/L). The experiments were carried out using the natural photoperiod and temperature conditions.

### Aquatic Environments and Water Provision

The experimental setup involved the selection of glass tanks with a capacity of 40 liters. These tanks were filled with tap water that had been treated to remove chlorine prior to the commencement of the experiment. The glass tank was subjected to a comprehensive cleaning process, ensuring its cleanliness. Tap water was then held for a duration of 48 hours, after which it was utilized for a complete refill of the tank in the morning. The quality of water was periodically assessed using established methodologies as outlined in the APHA (1998) standard guidelines.

### Chemical Substances and Laboratory Equipment

The present work utilized analytical grade chemicals and Borosil acid-alkali resistant glassware. All analytical procedures and the preparation of stock solutions were conducted using distilled water.

### Procedure for the preparation of the test solution

A stock solution of Chlorpyriphos (20% EC) was prepared and kept for further use. Various test dosages were created in order to create a dilution of the stock solution.

### Experimental Design

The fish were subjected to renewal bioassay in order to conduct short-term testing on acute toxicity over a period of 96 hours. The tests exclusively utilized fish specimens that exhibited robust health and demonstrated vigorous locomotion. A total of eight fish specimens were allocated to individual aquariums and subjected to varying doses of Chlorpyrifos, namely at levels of 0.4, 0.8, 1, 3, 5, 7, 10, 12, 15, and 19 parts per million (ppm). Control groups were maintained for each exposure. Each experimental session was conducted over a duration of 96 hours. The mortality rate of the fish was documented. The test media underwent daily renewal throughout the duration of the experiment. The physicochemical properties of the water, including temperature, pH, and dissolved oxygen, were regularly assessed using established protocols outlined in the APHA (1998) guidelines. To ensure repeatability, the impact of each concentration was assessed by a minimum of three repetitions. The data acquired during the course of the experiment were subjected to statistical analysis in order to determine if there is any correlation between various treatments (concentrations) and fish mortality (Bindu and Geetha, 2009). The LC_50_ values, together with their corresponding 95% confidence intervals, for various exposure durations were computed using the statistical software SPSS 24.

### Biochemical Investigations

Following a 96-hour exposure period, live fish specimens were euthanized, and specific tissues of interest, including muscle, intestine, and stomach, were carefully removed for the purpose of conducting biochemical analyses. The protein content of the stomach, intestine, and muscle was assessed using the Folin-Ciocalteu method. The determination of glycogen content in tissues was conducted using the Anthrone method as described by Seifter et al. (1959), following the methodology outlined by Lowry et al. (1951).

### Preparation of tissue for light microscopy studies

Following a 96-hour exposure period, the LC_50_ of Chlorpyriphos was determined for fishes over a four-day period. Subsequently, both control and experimental groups of fishes were sacrificed, and their muscle, intestine, and stomach were meticulously dissected. The dissected tissues were then carefully preserved and hardened using neutral buffered formalin (NBF) for a duration of two days. This preservation method facilitates chemical and physical alterations within the tissue by gradually permeating it. In order to induce dehydration, the tissues were subjected to a sequential immersion process involving alcohol solutions of progressively higher concentrations until pure, anhydrous alcohol was attained (at concentrations of 70%, 90%, and 100%). Subsequently, the tissues were cleaned using methyl benzoate. In order to facilitate infiltration, two cycles of wax immersion were conducted, with each cycle lasting for one hour at a temperature of 60°C. The tissues were fixed in paraffin and subsequently shaped into blocks. Tissue samples with a thickness of 5 μm were prepared and subjected to staining using hematoxylin and eosin. The stained sections were examined using a compound research microscope, and photomicrographs were captured using a photomicrography equipment.

### Statistical Analysis

The LC_50_ values of Chlorpyriphos were calculated at various exposure durations using the statistical software SPSS 24. The upper and lower confidence bounds were computed, along with the slope functions. In the context of biochemical research, the mean values were calculated using all replicates. Statistical analysis was conducted using the student’s test to examine the variations in biochemical parameters across different concentrations. The mean ± standard error was used to express the biochemical parameters of various tissues.

## III. RESULTS

Aquatic organisms are extremely vulnerable to the toxic effects of chemicals and insecticides since the aquatic medium is a very efficient solvent for chemicals and these toxicants enter the body of organisms via absorption or oral intake from the immediate environment. Fish have very inefficient mixed-function oxidase systems to detoxify the insecticides which makes them vulnerable to environmental contaminants. In the present study, a static acute toxicity test was conducted to determine the LC_50_ of Chlorpyrifos on A.testudineus for 24, 48, 72 and 96h upper and lower confidence limit and presented in Table.1. In control and lower concentrations (0.462, 0.187, 0.074ppm) mortality was nil throughout the experiment (96h) whereas, in higher concentrations like 15.61, 18.97, 20ppm mortality was observed before the completion of 24h. The median lethal concentration of Chlorpyrifos for 24 hours is 11.59 ppm and that of 96h is 3.82 ppm indicating that the mortality was positively correlated with an increase in concentrations of Chlorpyrifos as well as exposure period.

**Table 1.** Table showing the LC_50_ values, slope function, upper and lower confidence limits for Chlorpyrifos at different time intervals for the fish *Anabas testudineus*.

| Parameters | 24 hour | 48 hour | 72 hour | 96 hour |
| --- | --- | --- | --- | --- |
| LC<sub>50</sub> (ppm) | 11.59 | 8.82 | 6.30 | 3.82 |
| Slope function | 7.19 | 6.25 | 5.03 | 3.41 |
| Upper confidence limit | 13.67 | 10.48 | 7.75 | 5.11 |
| Lower confidence limit | 9.72 | 7.14 | 4.77 | 2.55 |

### Biochemical studies

In determining the physiological status of fish and recognizing acute or chronic toxicity of insecticides on target organs of fish biochemical tests are useful and routine laboratory tests. In our study, the protein content of the stomach, intestine and muscle of *A*.*testudineus* exposed to 96h LC_50_ for four days showed a significant decrease from the control. Estimation of glycogen content of the stomach, intestine and muscle of *A*.*testudineus* for the same exposure period also indicates a significant decrease in exposed tissues from control. The results are presented in Fig.2 and 3.

**Fig 1.**
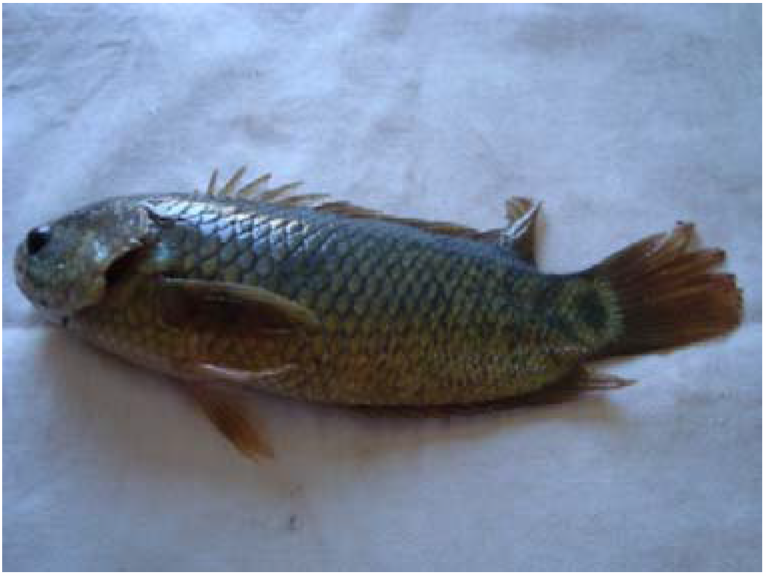
*Anabas testudineus*-test organism

**Fig 2.**
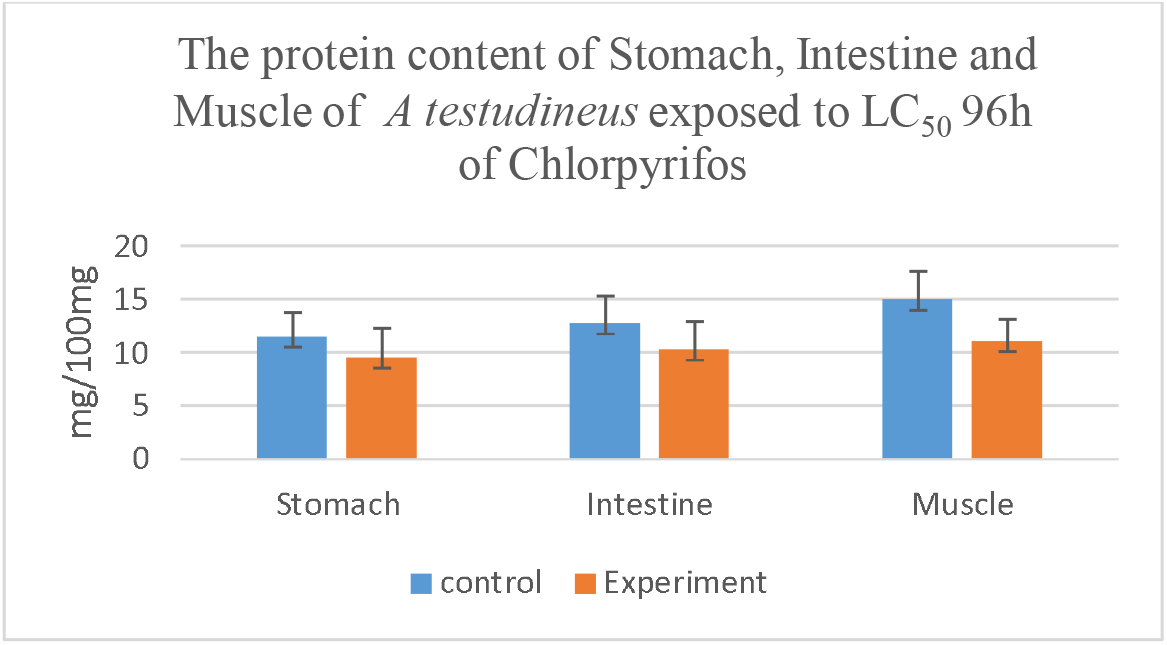
The protein content of the Stomach, Intestine and Muscle of *Anabas testudineus* exposed to LC_50_ of Chlorpyrifos for 96h.

**Fig 3.**
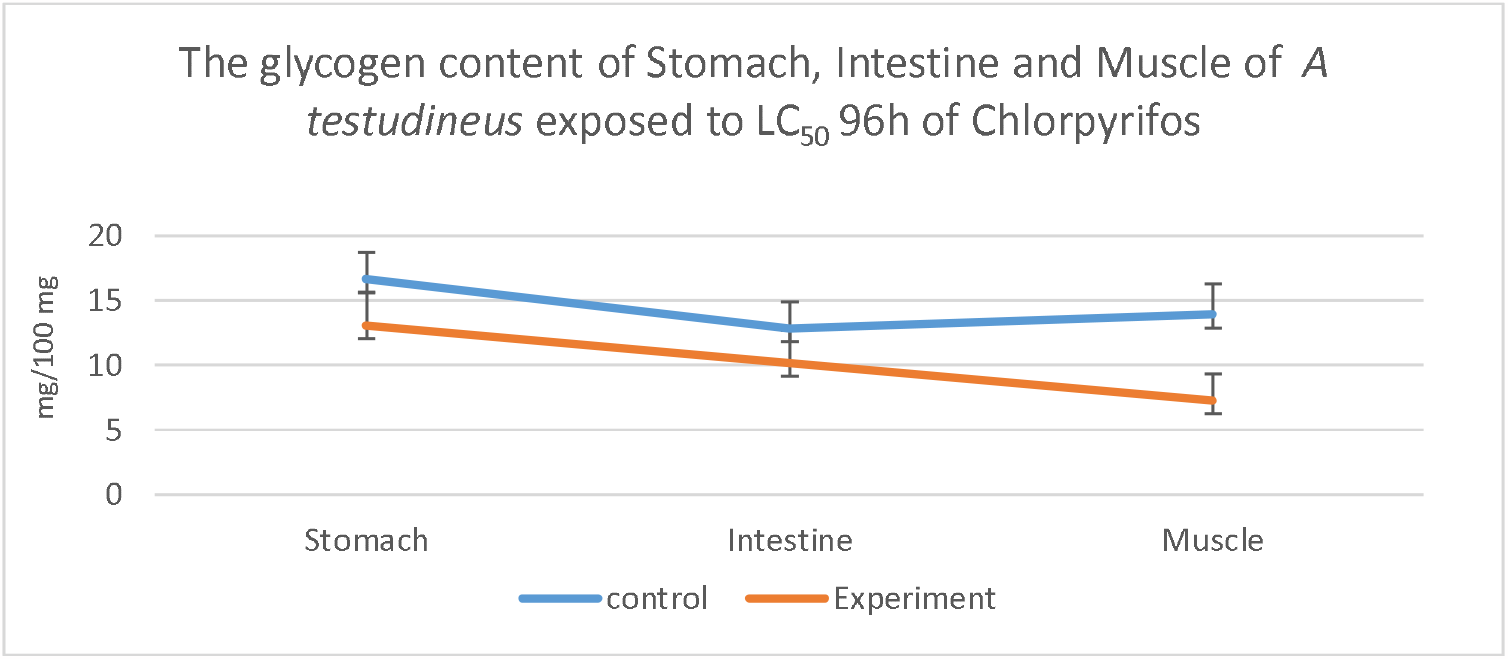
The glycogen content of the Stomach, Intestine and Muscle of *Anabas testudineus* exposed to LC_50_ of Chlorpyrifos for 96h.

### Histopathological Studies

Histopathological investigations provide information about the health and functionality of organs of fishes and help in monitoring water pollution since tissue injuries and damages results in reduced growth and fitness, decreased survival and low reproductive success of fishes and the intensity of damage depend on the concentrations of insecticides and the length of the period fish are exposed to toxicants. In the present study, histology of tissues (stomach, intestine and muscle) of *A*.*testudineus* exposed to 96h LC_50_ for 4 days showed significant alterations from the control structure

#### Stomach

The stomach with numerous folds in its lining is sigmoid and highly distensible. There are many layers of muscle, including a muscularis mucosa with adjacent layers of connective tissue often containing large numbers of eosinophilic granule cells. There are four main layers in the tube such as a serosa of connective tissue surrounded by simple squamous peritoneal epithelium, a muscular coat often divided into an inner circular and an outer longitudinal layer, a submucosa less cellular connective tissue layer with blood vessels, lymphatic tissue and nerve plexi and a mucosa, lining the lumen of the tube and consisting of an inner epithelium, a middle lamina propria (a cellular connective tissue) and outer muscularis mucosa; Stomach sections of *A. testudineus* exposed to 3.82 ppm (LC_50_96 hour) of Chlorpyrifos for 96 hours showed degeneration of the mucosa layer, necrosis of mucosa epithelium (Fig.5). Control sections of the stomach of *A. testudineus* show no histological changes (Fig.4).

**Fig 4.**
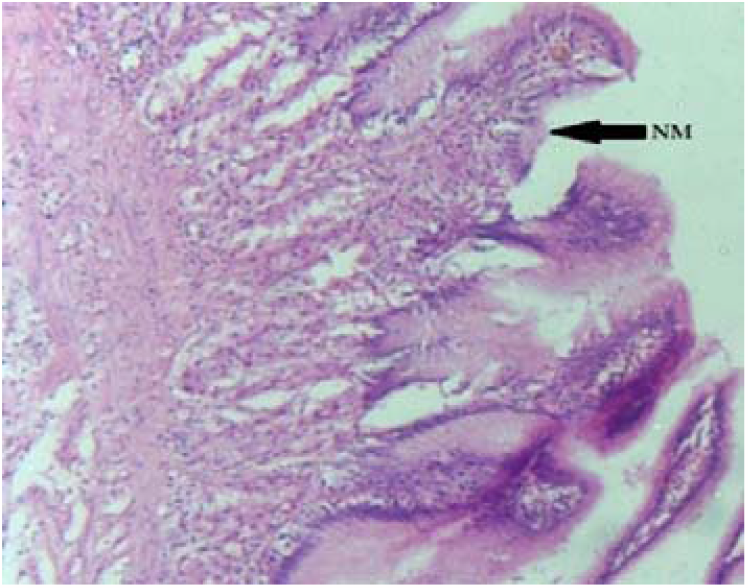
Control section of stomach showing normal mucosa

**Fig 5.**
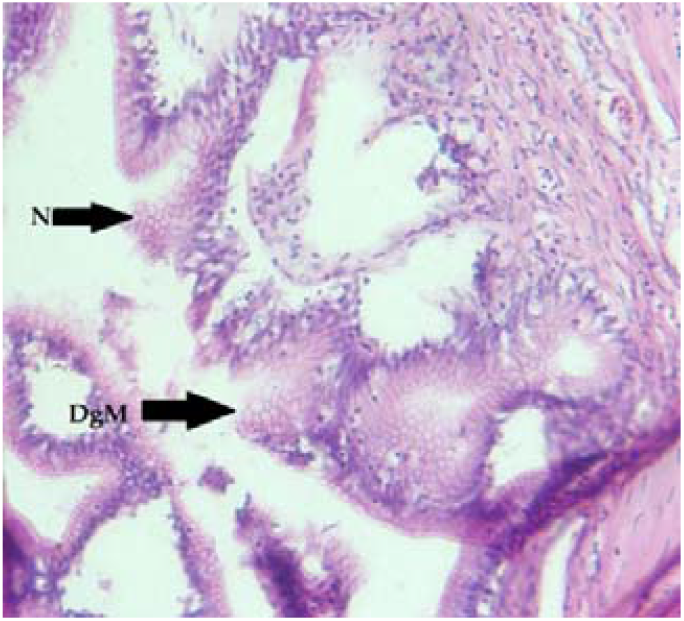
Cross section of stomach of treated fish showing degenerative mucosa (DgM) and necrosis of musoca epithelium (N)

### Intestine

The intestine of fish consisted of four basic layers-mucosa, submucosa, muscularis and serosa. The serosa was very thin and was made up of a single layer of peritoneal cells, and also had blood capillaries. The muscularis mucosa consisted of two layers. The outer one was the thin layer of longitudinal muscle fibers and the inner one was the thick layer of circular muscle fibers. The submucosa consisted of loose connective tissue fibers. The Mucosa layer was thrown into prominent folds forming villi and was made up of a single layer of simple columnar epithelium, which was formed of absorptive cells and goblet cells. The extension of the submucosa in the villi was the lamina propria. Villi consisted of absorptive and mucus-secreting cells.

Histology of the intestine of *A. testudineus* exposed to 3.82 ppm (LC_50_96 hour) of Chlorpyrifos for 96 hours showed partly disintegrated Serosa, muscularis layer was highly swollen and circular muscle layer damaged in some areas, necrosis in the Mucosa layer, Sub mucosa was extensively damaged and irregular (Fig.7). Control sections of the intestine of *A. testudineus* show no histological changes (Fig.6)

**Fig 6.**
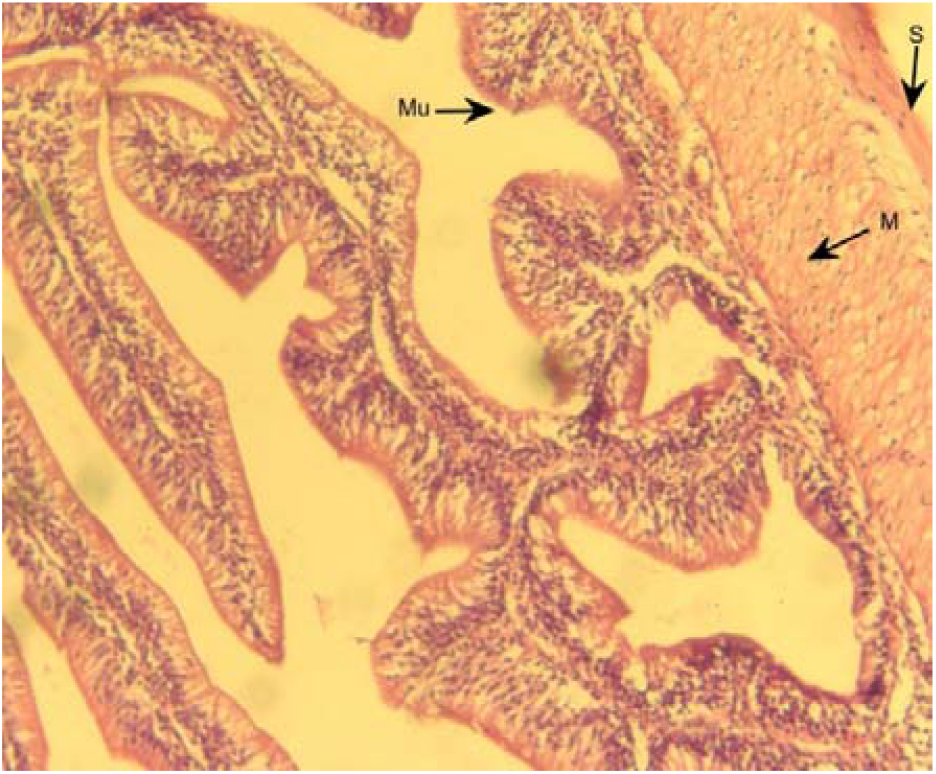
Control section of intestine showing normal serosa (S), mucosa (Mu) and muscular layer (M)

**Fig 7.**
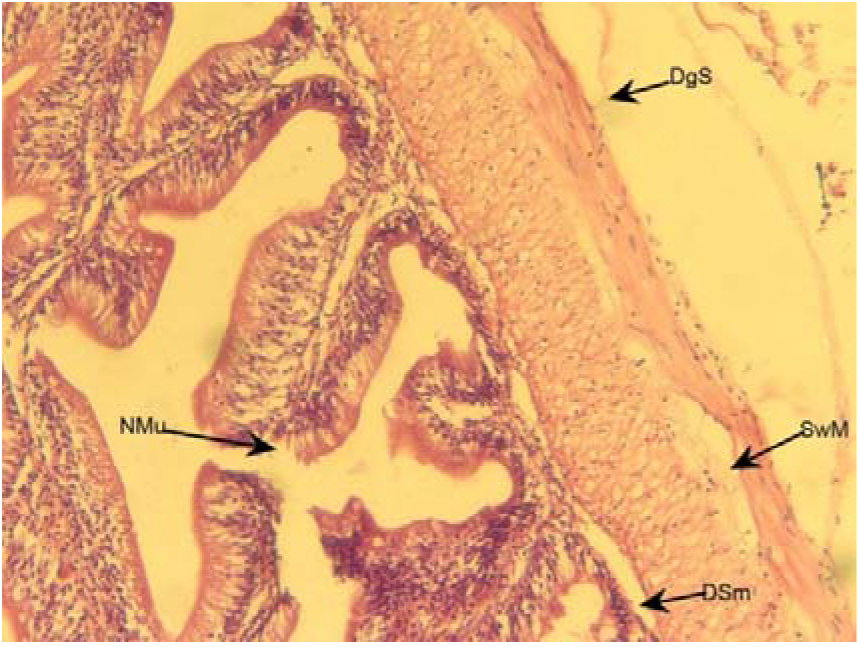
Cross section of intestine of treated fish showing degenerative serosa (DgS), damaged and irregular submucosa (DSm), necrosis of mucosa (NMu) and swelling of muscular layer (SwM)

### Skeletal muscle

The striated muscle cell of the skeletal muscles is multinucleated and in each cell, there are several longitudinal myofibrils, each of which is comprised of several myofilaments. There are red and white muscle fibers, for two kinds of swimming activity: the red fibers are related to sustained activity, while the white fibers are too short, strong bursts of motion. The greatest volume of body tissue comprises white muscle with low numbers of mitochondria and low respiratory activity. The white fibers are anaerobic, fast-contracting and fast-fatiguing fibers.

Histopathological alterations in the muscles of *A. testudineus* exposed to 3.82 ppm (LC_50_96 hour) of Chlorpyrifos for 96 hours included degeneration in muscle bundle accompanied with focal areas of necrosis and also, formation of vacuoles in muscle bundles (Fig.9). Control sections of the muscle of *A. testudineus* show no histological changes (Fig.8) DISCUSSION

**Fig 8.**
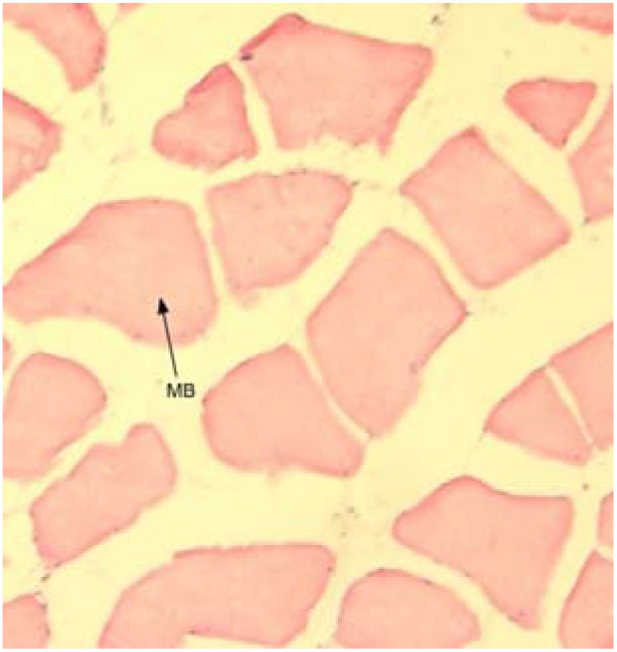
Control muscle showing normal muscle bundle

**Fig 9.**
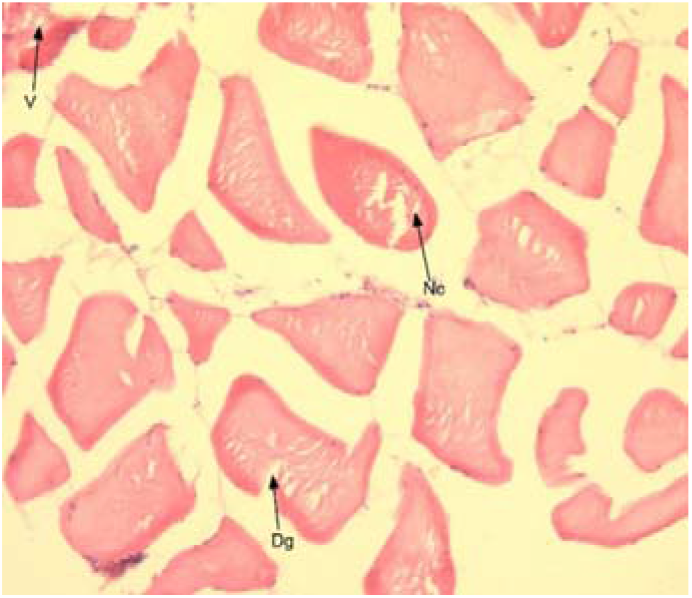
Section of muscle of treated fish showingdegeneration of muscle bundle (Dg), necrosis (Nc) and formation of vacuoles (V)

The LC_50_ values of Chlorpyrifos on *A. testudineus* at 24, 48, 72, and 96 hours were determined to be 11.59, 8.82, 6.30, and 3.82 parts per million (ppm), respectively. The death of fish exhibits a direct association with the concentration of the toxicant, as observed through an increase in mortality with higher concentrations. The LC_50_ value of dimethoate on Clarias batrachus, a species of catfish, was reported to be 65 mg/L by Begum and Vijayaraghavan in 1995. In a study conducted by Dikshith and Raizada in 1981, the 96-hour LC_50_ value for dimethoate on Channa punctatus, a type of air-breathing teleost, was reported as 47 mg/L. According to Usha et al. (2017), the 96-hour LC_50_ value for Chlorpyrifos on Oreochromis mossambicus was determined to be 29.99 mg/L. According to a study conducted by Verma and Rawat in 2017, the LC_50_ value of Chlorpyrifos on Heteropneustes fossilis after a 96-hour exposure period was determined to be 4.64 mg/L. The behavioral and physiological systems of the organisms, specifically the fish, are impacted by any chemical modifications in a natural aquatic environment (Radhaiah et al., 1987). According to Kegley et al. (1999), even pesticide concentrations that do not reach lethal levels for fish can nevertheless have notable effects on their behavior and physiology, ultimately leading to impairments in both survival and reproductive capabilities. The current investigation observed a notable reduction in the protein content of the muscle, stomach, and intestine of fish that were exposed to the 96-hour LC_50_ concentration of Chlorpyrifos for a duration of four days, when compared to the protein content of fish in the control group. Singh et al. (2010), Martin et al. (2008), Waghmare and Wani (2014), Patil and David (2009), and Ramesh and Saravanan (2008) have reported a decrease in protein content in Colisa fasciatus when exposed to cypermethrin, Catla catla when exposed to mercury chloride, Labeo rohita when exposed to polo and malathion, and Cyprinis carpio when exposed to chlorpyrifos. According to Umminger (1977), chlorpyrifos induces hyperactivity as a response to stress, hence meeting the increased energy requirement. This heightened activity subsequently leads to the depletion of food reserves. The observed reduction in protein levels upon exposure to chlorpyrifos may indicate the potential degradation of proteins as a means for organisms to obtain energy (Tulsi and Jayantha, 2013). When organisms are subjected to toxicant-induced stress, they require additional energy to counteract the stress, and proteins can serve as an energy source for fish. Consequently, when there is insufficient energy from alternative sources such as carbohydrates, protein depletion occurs in the tissues (Neff, 1985). The glycogen levels in the stomach, gut, and muscle of *A. testudineus* exhibited a notable reduction after being subjected to a four-day exposure of Chlorpyrifos at its 96-hour LC_50_ concentration. This observation suggests that the toxicant has enduring and cumulative inhibitory impacts on glycogen content. The depletion of glucose in specific tissues can be attributed to its rapid utilization in order to meet the energy requirements during toxic manifestations. Additionally, this depletion may also be influenced by the differential accumulation and distribution of toxic substances in various tissues, which is contingent upon factors such as age, temperature, perfusion, vascularity, and residual blood volume (Villarrel and Navorro, 1987).

The frequency and severity of tissue lesions in fish are influenced by the concentrations of insecticides and the duration of exposure to these poisons. It has been observed that several pesticides can cause histopathological damage in fish, either through specific or non-specific mechanisms (Fanta et al., 2003). In our study, it was observed that exposure of *A. testudineus* to the 96 h LC_50_ of Chlorpyrifos for a duration of four days resulted in the degeneration of the mucosa layer and necrosis of the mucosa epithelium in the stomach. Additionally, the serosa showed partial disintegration, the muscularis layer was highly swollen, and the circular muscle layer was damaged. Furthermore, the mucosa layer exhibited necrosis, and the submucosa of the intestine was extensively damaged and irregular. These observed pathological changes may have disrupted the normal processes of digestion and absorption in the affected organisms. The absorptive and secretory capabilities of fish were impaired due to the exposure to several toxicants, resulting in necrosis of epithelial cells, degeneration of epithelium and sub-epithelial connective tissue, and the presence of lesions on intestinal villi (Schiller, 1979).The development of cell lysis, characterized by the rupture of cells and the removal of the cell nucleus, is closely linked to necrosis, as indicated by Sulastri et al. (2018). The phenomenon of cell degeneration is known to result in the modification or disturbance of cellular functionality, as documented by Rahayu et al. (2013).According to Muley et al. (1996), the exposure of Channa punctatus to carbofuran resulted in the deterioration of the mucosal folds’ structural integrity, as well as the degeneration and necrosis of the submucosa in the gut. Additionally, the study observed degenerative alterations and rupture in the tip of the villi. According to Srivastava and Srivastava (1995), the exposure of Heteropneustes fossilis to Selenium resulted in the fusion of submucosa with muscles, the loss of columnar epithelium, and the disruption of serosa. The histopathological changes observed in the muscles of *A. testudineus* following a 96-hour exposure to Chlorpyrifos at a concentration of 3.82 ppm (LC_50_ for 96 hours) consist of degeneration in muscle bundles, accompanied by focal areas of necrosis. Additionally, the creation of vacuoles inside the muscle bundles is also evident. The histopathological changes observed in the muscle tissue of *A. testudineus* are consistent with the histological alterations documented in other fish species, such as the African catfish Clarias gariepinus (Islam et al., 2021), Hoplias alabaricus (Miranda et al., 2008), Mugil capito red belly Tilapia, Tilapia zillii (Mohamed and Gad, 2008), Solea vulgaris (Mohamed, 2009), and zebrafish, Danio rerio (Bhuvaneshwari, 2015), following exposure to various insecticides.

## IV. CONCLUSIONS

The current investigation examined the toxicity of chlorpyrifos on *A. testudineus* and observed a positive correlation between toxicity and both concentration and exposure duration. The presence of histopathological changes in the stomach, gut, and muscle of the fish may have an indirect impact on individual survival and reproductive success, thereby disrupting the population dynamics of A. testudineus. The extensive and unselective application of this pesticide in agricultural settings, particularly in close proximity to aquatic environments, poses significant risks to aquatic creatures and has the potential to impact several trophic levels within the food chain, ultimately resulting in a substantial disturbance to aquatic fauna.

## AUTHORS’ CONTRIBUTION

All authors equally participated in the design and conduct of the experiment, data analysis and preparation of the manuscript.

## DATA AVAILABILITY STATEMENT

The data that support the findings of this study are available at a reasonable request from the corresponding author.

